# Probiotic Stirred Yogurt -Mediated the Prevention of Metabolic Dysfunction-Associated Steatotic Liver Disease in Rats

**DOI:** 10.1101/2024.11.23.624973

**Authors:** Mohamed S. Sallam, Eman H. Ayad, Saeid M. Darwish, Mohamed A. Gomaa, Marwa G. Allam

**Affiliations:** Department of Food Science, Faculty of Agriculture, Saba Basha, Alexandria University, Alexandria, Egypt 3

**Keywords:** Probiotics, Non-alcoholic fatty liver disease (NAFLD), Lipid profile, Liver Histology, Glucose, High-fat diet (HFD)

## Abstract

Metabolic dysfunction-associated Steatotic liver disease (MASLD) is the most common cause of chronic liver diseases and it is greatly influenced by an unhealthy and sedentary lifestyle. Studies have encouraged the use of probiotics as promising and safe therapeutic approach. This research examined the probiotic potential of eleven bacterial strains, with *Lactococcus rhamnosus*, L*actobacillus plantarum, Lactobacillus acidophilus*, and *Bifidobacterium lactis* being selected for yogurt production, either as single strain or in combination. The probiotic-rich yogurt samples were then tested for their ability to prevent the development of MASLD in *vivo* using a rat model fed a high-fat diet (HFD). Key biomarkers such as cholesterol, triglycerides, low-density lipoproteins (LDL), high-density lipoproteins (HDL), glucose, insulin, alanine transaminase (ALT), and albumin levels were measured. Besides, liver histology and the microbial analysis of the small intestine were examined. The sensory revealed that yogurt containing the probiotic mixture (T5) had improved taste and flavor. The probiotic mixing revealed a potentiated effect in preventing MASLD. T5 significantly lowered cholesterol (85.1 mg/dl), triglycerides (122.3 mg/dl), LDL (25.8 mg/dl), glucose (28.6 mg/dl), insulin (0.84 μIU/ml), and ALT (36.9 U/L) levels in the rats to close values as in the control. Additionally, the treated rats showed restored albumin and HDL levels compared to those treated with yogurt containing a single probiotic strain. The ameliorating effects of T5 were also evident in the liver histology, with normal hepatocytes and minimal fat deposits. Furthermore, the probiotic mixture in T5 improved the liver-gut axis by reducing the coliform and *staphylococcus* count in the small intestine. Consequently, the probiotic mixture could be a promising therapeutic agent for MASLD and the ultimate goal of this project is to develop a functional food product that can provide an alternative cost-effective approach for the management of MASLD in Egyptian population.

## 1. Introduction

Metabolic dysfunction-associated Steatotic liver disease (MASLD) previously termed non-alcoholic fatty liver disease (NAFLD), is the most common cause of chronic liver disease, affecting 24% of the world population with increasing global prevalence (**Younossi, Anstee et al. 2018)** Steatosis, the first stage, is characterized by the deposition of triglycerides within the hepatocytes, which is evidenced by more than 5% of hepatocytes containing visible lipid droplets, in the absence of other compromising factors such as viral infection, chronic liver disease, or excessive alcohol consumption (**Buzzetti, Pinzani et al. 2016)**. This stage is relatively benign and has not been linked to significant changes in liver function. However, steatosis may progress to metabolic dysfunction-associated steatohepatitis (MASH) in nearly 30% of the cases, which is characterized by inflammation and hepatocellular ballooning, with a potential progression to fibrosis in 40% of MASH patients **(Younossi, Koenig et al. 2016)**. Moreover, the disease progression involves inflammatory markers that lead to further hepatocellular lesions, including necrosis and apoptosis via hepatic stellate cell-mediated signaling **(Khomich, Ivanov et al. 2019)**. Generally, MASLD affects about 24% of the world’s population, with specific geographical and racial prevalence rates. South America and the Middle East were the most affected regions (31 and 32%, respectively), followed by Asia (27%), north America (24%), Europe (24%), and Africa (13%). It was also estimated that the global prevalence of MASH was between 1.5 and 6.45%, with more than half (59.1%) of MASLD patients affected. Moreover, 9% of MASH patients were shown to progress to advanced fibrosis; follow-up studies revealed an estimated 2.6% annual incidence of MASH-related cirrhosis and 52.9% incidence of hepatocellular carcinoma in MASH patients **(Younossi, Koenig et al. 2016)** Therefore, ameliorating liver diseases has become vital. Dietary patterns and healthy food are among the common, successful approaches to alleviating MASLD (**Montemayor, García et al. 2023)**

Fermented dairy products show positive management in MASLD patients by reducing body weight and improving insulin sensitivity through the modification of the gut microbiota **(Mackowiak 2013, Montemayor, García et al. 2023)** Yogurt is one of the most popular fermented dairy products worldwide, which is usually manufactured by fermented milk with *Streptococcus thermophiles* and *Lactobacillus delbrueckii* ssp. *Bulgaricus* **(Illikoud, Mantel et al. 2022)**. However, neither species survives well during the passage through the gastrointestinal tract, reducing their ability to confer probiotic benefits **(Verma, Patel et al. 2022)**. Thus, fortifying the yogurt with different, robust probiotic strains could show a more positive, probiotic influence.

Probiotics, according to the World Health Organization, are “Live microorganisms, which when administered in adequate amounts confer a health benefit on the host” (**Hill, Guarner et al. 2014)**. Probiotics, particularly Lactobacillus and Bifidobacterium, can restore the microbiota balance in the gut, positively influencing the gut-liver axis. It has shown a potentiated reduction in fat accumulation in the liver, inflammation, and oxidative liver damage (**Montemayor, García et al. 2023)** Therefore, combining the benefits of yogurt and probiotics could show a potential MASLD management. Commercial yogurt is usually made with the two organisms, Streptococcus thermophiles and Lactobacillus delbrueckii ssp. bulgaricus from cow’s milk. The two organisms complement each other in a synergistic manner **(Illikoud, Mantel et al. 2022)**.However, neither species survives well during the passage through the gastrointestinal Consequently, it is unlikely that they are able to confer probiotic benefits. Therefore the present study aims to assess the probiotic potential of different bacterial strains and fortify yogurt samples with these strains. The acceptability of the probiotic yogurt was evaluated and tested for its ability to prevent MASLD in a high-fat diet rat model.

## 2. Materials and Methods

### 2.1 Materials

#### Bacterial strains

The Probiotics strains used in this experiment was *lactobacillus rhamnosus* (LGG), *Lactobacillus plantarum* (Lp-115), *and lactobacillus acidophilus* (La-5) and *Bifidobacterium lactis* (BL-04), *Lactobacillus salivarius* (Ls-33), *Bifidobacterium breve* (Bb-03), *Bifidobacterium longum* (Bl-05), *lactobacillus casei* (Lc-11), *Lactobacillus fermentum* (Lf-10) *lactobacillus reuteri* (Lr-10) and *Lactobacillus paracasei* (Lp-33). In addition to the yoghurt commercial starter culture (YOFLEX®), consisting of (*Streptococcus Thermophilus* and *Lactobacillus delbrueckii* subsp. *bulgaricus*) all bacterial strains was obtained from (Chr. Hansen; Denmark) and were stored at -18°C until further use.

#### Media and Media constituents

De Mann, Rogosa and Sharpe (MRS) broth, M17 broth, Staph 110 media and violet red Bile Agar (VRBA) were obtained from Bio Life (Italy). Plat count Agar (PCA) and Rose Bengal media were obtained from Oxoid (England).

### 2.2 Preparation of probiotics culture

Cultures were prepared according to the culture collection’s recommendation. The lyophilized pellet of each strain were activated in 10 ml of sterilized MRS broth and incubated at 37 ºC for 24 h. Then, the activated cultures were purified on MRS agar; the agar plates were incubated at the recommended temperature until colonies appeared (48 h). One of the colonies was transferred to 10 ml MRS broth media and incubated at the recommended temperature for 24 h. The purity of the cultures was checked by catalase test and microscopic examination after Gram staining. The pure confirmed culture (1 ml) was transferred into a sterile 1.5 ml Eppendorf containing 200 μl of sterile glycerol and stored at -18 ºC. Prior to use, strains were subcultured twice in MRS broth (10 % v/v) and incubated aerobically for 24 h at 37 ºC or 28 ºC.

### 2.3 Assessment of potential probiotic strains

#### 2.3.1 Acid resistance

Overnight cultures were previously prepared by inoculating the strains into MRS broth (0.1% v/v) at 37°C for 16 h to reach the log phase. Then, MRS broth was acidified using HCL to obtain media with different pH values of 2, 3, and 4. Subsequently, the acidified media was mixed with the bacterial cultures at a ratio of 10% (v/v). The mixtures were incubated at 37°C and the growth was followed for 6 h by measuring the optical density at 650nm (O.D650) using a spectrophotometer (Apel-pD-303UV Spectrophotometer, Japan). The growth of the cultures in each broth was compared to the standard MRS broth (pH 6.6) as described by Baccigalupi, Di Donato et al. (2005)

#### 2.3.2 Bile salts resistance

The strains were evaluated for growth rapidity in MRS broth media with and without bile salts. MRS broth was prepared by adding bile salts (Bio Life, Milano, Italy) to the media at concentrations of 0, 0.2, 0.3, and 0.4% (w/v). Afterward, overnight-grown cultures were inoculated into the media and incubated at 37°C. Bacterial growth was monitored over 6 h by measuring the optical density at 650 nm (OD650). The comparison of cultures was based on their growth rates in each broth as described by Baccigalupi, Di Donato et al. (2005)

#### 2.3.3 Phenol tolerance

The growth ability of the strains in the presence of phenol was evaluated according to the method described by Aswathy, Ismail et al. (2008) MRS broth-activated cultures (24 hours old) were adjusted to an absorbance of 0.8 at 600 nm. The cultures were then inoculated into MRS broth media containing 0, 0.2, and 0.4% phenol at 1% (v/v) and incubated at 37 ºC for 24 h. The growth was assessed by measuring the absorbance at 600 nm after 24 h.

### 2.4 Manufacture of stirred yoghurt

Cow milk was standardized using skimmed milk (SMP) to a solids-non-fat content of 13%. The milk was heat-treated at 85ºC/15 min. Afterward, the milk was cooled to 42±1 ºC, divided into six equal portions, and inoculated with 0.01 g/kg the following probiotic strains: *Lactococcus rhamnosus* (T1), L*actobacillus plantarum* (T2), *Lactobacillus acidophilus* (T3), *Bifidobacterium lactis* (T4), and the mixture of the four mentioned probiotics (1:1:1:1) (T5). After an hour all samples were inoculated with the commercial yogurt starter culture (0.03g/kg). A control group (T0) was also prepared by adding only the yogurt starter culture (YOFLEX®). The inoculated milk samples were then poured into 100-ml plastic cups and incubated at 42±1ºC until set coagulation at pH ∼4.6 (about 5 h.). The samples were then left to cool down and stored at 4°C until further use.

### 2.5 Sensory evaluation

The sensory attributes of the fresh probiotic-stirred yogurt samples were evaluated by 20 consumer-oriented panelists (11 men and 9 women) at the Food Science Department, Faculty of Agriculture, Alexandria University as described by **(Darwish, Soliman et al. 2021)**, to evaluate 6 groups, T0 group contains the commercial starter culture, T1group contains commercial starter culture plus *lactobacillus Rhamnosus*, T2 group contains commercial starter culture plus *lactobacillus Plantarum*, T3 contains commercial starter culture plus *lactobacillus acidophilus*, T4 group contains commercial starter culture plus *Bifidobacterium lactis* and T5 group contains commercial starter culture plus mixture of the four mentioned probiotic bacteria. The criteria for selection depended on their experience and background related to the fermented dairy products. The representative samples which had not damaged or changed during cold storage at 4° C, were allowed to rest at room temperature 25° C for 10 min before evaluation. panellists were allowed to evaluate the samples independently using a 7-point Hedonic scale. This scale consisted of the test parameters of color, taste, flavor, odor, texture, appearance, acidity and overall acceptability, accompanied by a scale of seven categories as: 1 = dislike extremely; 2 = dislike much; 3 = dislike slightly, 4 = neither dislike nor like, 5 = like slightly; 6= like much; 7 = like extremely. Panellists were instructed to rinse their mouth with water between samples to eliminate any residual effects.

### 2.6 Feeding experiment

#### 2.6.1 Animals diet

The commercial chow, used as a normal diet for rats, was obtained from the Home Economics Department, Faculty of Agriculture Alexandria University, Egypt. The normal diet is composed of soybeans (11.50 gm.), corn (5 gm.), starch (43.50 gm.), sucrose (30 gm.), salt mixture (5 gm.), cellulose (1 gm.), and vitamins mixture (1 gm.). Simultaneously, a high-fat diet (HFD) was prepared by adding 35% palm oil to the normal chow diet to increase the total fat content and energy (Table 1).

**Table (1):**
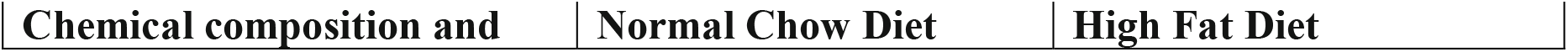
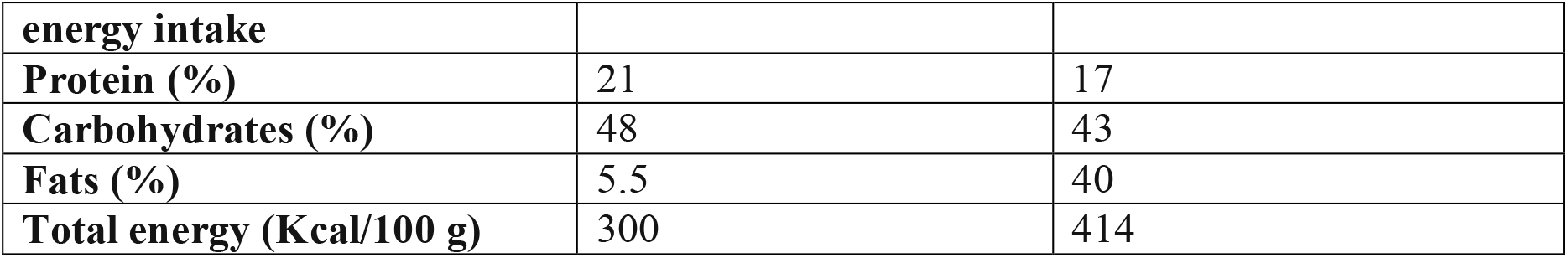
Proximate composition and total energy of Chow diet and High-fat diet.

Forty-eight male albino rats (4 weeks old; 68±1.5 gm.) were obtained from the Institute of Graduate Studies, Alexandria University, Egypt. The rats were housed in cages at 20±4°C with 40% humidity and a 12 h light/dark cycle. The rats were acclimatized for 7 days before starting the experiment, with free access to water or and the chow diet

#### 2.6.2 Experiment design

The rats were divided into eight groups, each consisting of six rats. During the first two weeks, all groups were fed the chow diet while receiving different treatments. The treatments were introduced daily by the gavage as follows: Groups I (negative control) and II (positive control) received water; while Groups III, IV, V, VI, VII, and VIII received yogurt samples T0, T1, T2, T3, T4, and T5, respectively (Table 2). After the initial two weeks, the chow diet was replaced with the high-fat diet in groups III to VIII, and they continued receiving their respective treatments. The weight of the rats was recorded weekly; blood was collected every two weeks from the medial canthus of the eye, using a micro hematocrit capillary tube during the fasting period (8 h). After 8 weeks, the rats were fasted for 8 hours, anesthetized, and sacrificed. Blood was collected for biochemical measurements; the organs (liver, kidney, and spleen) were washed with cold saline, dried, and weighed to determine the percentage of each organ to the body mass. Furthermore, the small intestines were obtained for microbiological analysis. Subsequently, portions of the liver were fixed with 10% formalin for histological examination.

**Table (2):** Rat’s group and their daily routine.

|  |  |
| --- | --- |
| **Group I** | Fed on chow diet. |
| **Group II** | Fed on high Fat diet. |
| **Group III** | Fed on HFD and control yogurt (T0) |
| **Group IV** | Fed on HFD and T1 yogurt, containing *Lactococcus rhamnosus* |
| **Group V** | Fed on HFD and of T2 yogurt, containing *Lactobacillus plantarum* |
| **Group VI** | Fed on HFD and of T3 yogurt, containing *Lactobacillus acidophilus* |
| **Group VII** | Fed on HFD and T4 yogurt, containing *Bifidobacterium lactis* |
| **Group VII** | Fed on HFD and T5 yogurt, containing the mixture of the four probiotics |

#### 2.6.3 Blood analysis

To collect the serum, blood samples were allowed to clot in dry glass centrifuge tubes at 24 ± 3°C, then centrifuged at 1400 g for 20 min. The clear, non-hemolysis supernatant serum was collected using a clean, dry disposable plastic syringe, and was kept at −80 °C for subsequent biochemical measurements. Blood samples were analyzed for lipid profiles, including triglyceride (TG), total cholesterol (TC), and high-density lipoproteins (HDL). The low-density lipoprotein (LDL) level was calculated using the equation introduced by Friedewald, Levy et al. (1972) as follows:

LDL (mg/dl) = TC−HDL− (TG/5). In addition, alanine aminotransferase (ALT) and albumin were measured as indicators of hepatic function. Glucose and insulin levels were also determined in the serum as indicators of metabolic function. All measurements were performed following the instructions provided in the Vitro scient kit.

#### 2.6.4 Microbial analysis of small Intestine

The small intestines of rats from groups I, II, and VIII were washed with 20 ml of sterilized saline using a sterilized syringe into a sterilized flask. Serial dilutions were then prepared, and samples were immediately analyzed using the aseptic sterile dilution technique described by Klaver, Kingma et al. (1993)

The total bacterial count was determined using the conventional diluting pouring plate technique on PCA media, and the results were expressed as colony-forming units (CFU g-1). To enumerate the total coliforms and *staphylococci*, the dilutions were grown on the VRBA media (**Marshall 1993)** and Staph 110 media (**Khan, Zinnah et al. 2008)**, respectively.

#### 2.6.5 Histological analysis

The fixed liver portions were dehydrated in ascending concentrations of ethanol, cleared in xylene, and then embedded in paraffin wax. Tissue sections were cut into a thickness of 3-5 Microns and stained with hematoxylin and eosin (H&E) following the method of Bancroft and Stevens (1996). The stained tissues were then examined under light microscopy for histopathology evaluation.

### 2.7 Statistical analysis

The statistical analysis was performed using the IBM SPSS Statistics software, version 23.0 (IBM Corp, 2015, Armonk, NY). The data were analyzed using a one-way analysis of variance (ANOVA), followed by Duncan’s test. The results were presented as mean values of two replicates ± standard deviation, with a confidence level of 95% (*p <* 0.05).

## 3. Results

### 3.1. The probiotic potential of different lactic acid bacterial strains

The viability and activity of probiotics are essential in the lower digestive tract, so these organisms should withstand the adverse conditions encountered in the host’s upper gastrointestinal tract (GIT). However, many probiotics lack the ability to survive the harsh acidity and bile concentration commonly encountered in the GIT. Therefore, acid resistances in the stomach and bile tolerance in the small intestine are selective characteristics of effective probiotics. During the screening of eleven strains of lactic acid bacteria, *Lactobacillus rhamnosus* (LGG), *Lactobacillus plantarum* (Lp-115), *Lactobacillus acidophilus* (La-5), and *Bifidobacterium lactis* (BL-04) showed tolerance to acid, bile, and phenol (Table 3). Based on these results, they were selected for the production of stirred yogurt.

**Table (3):** Acid tolerance, bile salts tolerance and phenol tolerance of the probiotic bacteria strains.

|  |  |  |  |  |  |  |  |  |
| --- | --- | --- | --- | --- | --- | --- | --- | --- |
| **Probiotic strains** | **Acid tolerance (PH)** |  |  | **Bile salts tolerance (%)** |  |  | **Phenol tolerance (%)** |  |
|  | **2** | **3** | **4** | **0.2** | **0.3** | **0.4** | **0.2** | **0.4** |
| *Lactobacillus acidophilus* (La-05) | + | + | + | + | + | + | + | + |
| *Bifidobacterium lactis* (Bl-04) | - | + | + | + | + | + | + | + |
| *Lactobacillus rhamnosus* (LGG) | + | + | + | + | + | + | + | + |
| *Lactobacillus plantarum* (Lp-115) | + | + | + | + | + | + | + | + |
| *Bifidobacterium longum* (Bl-05) | - | + | + | + | + | - | + | + |
| *Bifidobacterium breve* (Bl-03) | - | - | - | + | - | - | - | - |
| *Lactobacillus casei* (Lc-11) | - | + | + | + | + | + | + | - |
| *Lactobacillus salivarius* (Ls-33) | - | - | + | + | + | + | + | + |
| *Lactobacillus fermentum* (Lf-10) | - | + | + | + | + | + | + | + |
| *Lactobacillus reuteri* (Lr-1) | + | + | + | + | + | - | + | - |
| *Lactobacillus paracasei* (Lp-33) | - | + | + | + | + | - | + | + |

Data are represented in means of (-) not detected

### 3.2. Sensory attributes of probiotic-rich yogurt

The sensory evaluation of the probiotic yogurt samples and the control is shown in Figure 1. The results indicate that adding a mixture of probiotic strains *Lactobacillus rhamnosus* (LGG), *Lactobacillus plantarum* (Lp-115), *Lactobacillus acidophilus* (La-5), and *Bifidobacterium lactis* (BL-04) to the yogurt starter culture enhanced the organoleptic properties of the yogurt compared to the other samples. T5 exhibited the best sensory attributes, particularly taste and texture.

**Figure 1:**
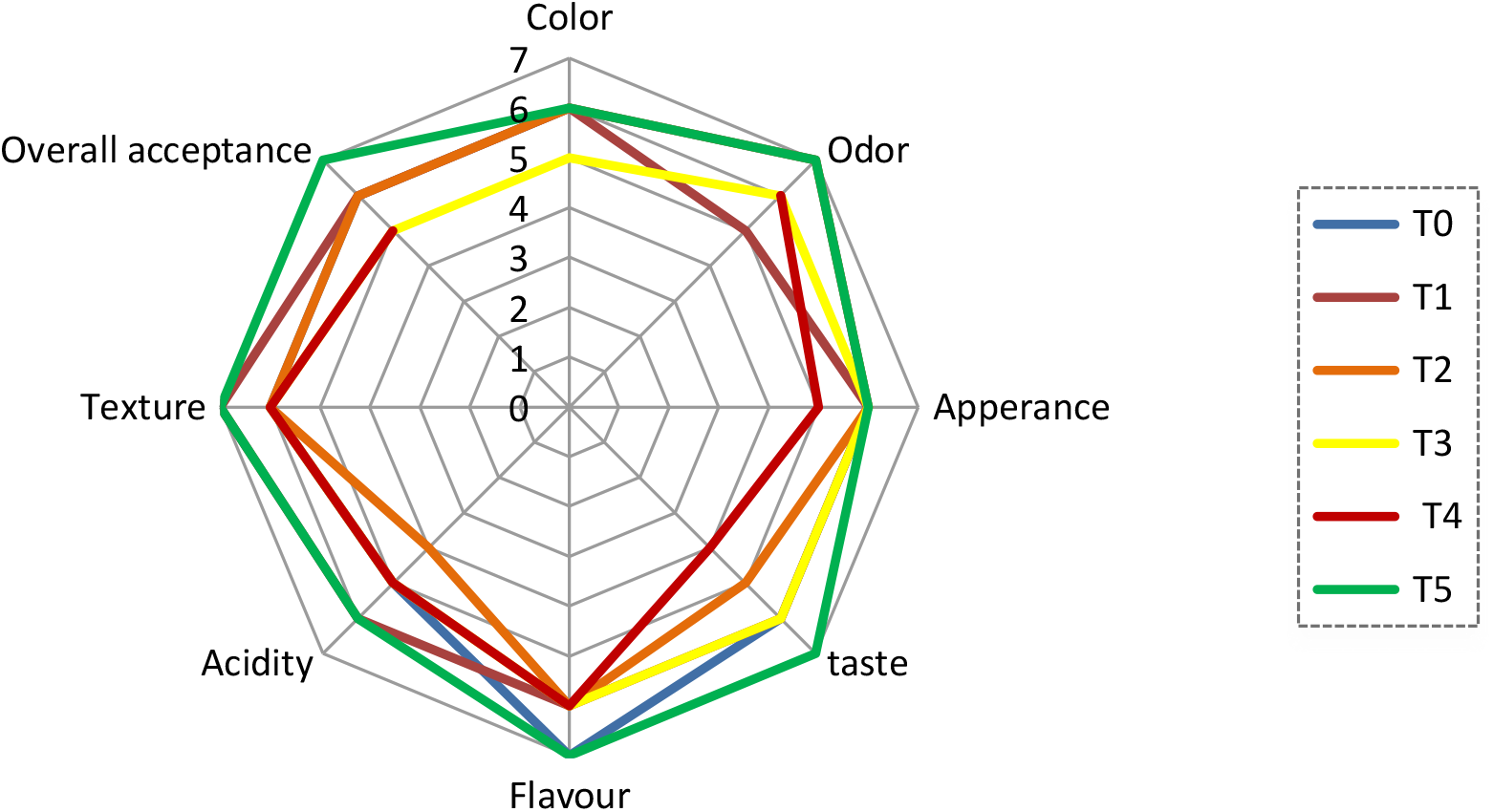
Sensory evaluation of yogurt samples enriched with different probiotic strains.

### 3.3. Effect of probiotics on the body weight of the rats

At the beginning of the experiment, there was no significant difference in body weight between the groups (p>0.05) (Table 4). However, while proceeding with the experiment, the final body weight and weight gain showed a significant decrease in the probiotics treated groups (Groups IV-VIII) compared to the positive control (Group II) that shows a significant increase in both. The probiotic treated groups had lower body weight and weights gain than the untreated Group II and group III. However, the inclusion of probiotics in Groups IV-VIII did not significantly reduce the body weight of the rats compared control negative Group III.

**Table (4):** Effect of probiotics on the body weight of the rats.

| Table (4): Effect of probiotics on the body weight of the rats |  |  |  |
| --- | --- | --- | --- |
|  | Initial body weight (g) | Final body weight (g) | Mean Body weight gain (g) |
| **Group I** | 66.2± 1.9<sup>a</sup> | 214.0± 3.7<sup>a</sup> | 147.8±3.7<sup>a</sup> |
| **Group II** | 66.0± 2.0<sup>a</sup> | 276.0±4.5<sup>c</sup> | 209.8±4.5<sup>c</sup> |
| **Group III** | 66.3± 1.9<sup>a</sup> | 249.0± 2.9<sup>b</sup> | 182.7±2.9<sup>b</sup> |
| **Group IV** | 67.1± 1.7<sup>a</sup> | 238.0±4.7<sup>b</sup> | 170.9±4.7<sup>b</sup> |
| **Group V** | 68.3± 1.6<sup>a</sup> | 244.0±2.8<sup>b</sup> | 175.7±2.8<sup>b</sup> |
| **Group VI** | 67.7± 1.7<sup>a</sup> | 239.0±3.5<sup>b</sup> | 171.3±3.5<sup>b</sup> |
| **Group VII** | 66.8± 1.9<sup>a</sup> | 239.0±4.1<sup>b</sup> | 172.2±4.1<sup>b</sup> |
| **Group VIII** | 66.7±1.9<sup>a</sup> | 231.0± 3.1<sup>b</sup> | 164.3± 3.1<sup>b</sup> |
| Data are presented as mean ± SD;<br>Means with different lowercase letters in the same column are significantly different (p<0.05). |  |  |  |

### 3.4. Effect of probiotics on the organs weight of the rats

In Group II, the HFD resulted in a significant increase in organ weight compared to the other groups (Table 5). The liver, spleen, and kidney increased by nearly 85%, 59%, and 31%, respectively, compared to the negative control. Both yogurt and probiotic-rich yogurt demonstrated an ameliorating effect on the organs of the rats, with the treated groups showing significantly lower organ weights than Group II. Notably, Group VIII, treated with T5 (a mixture of probiotics yogurt), mitigated the effect of the HFD on the liver, spleen, and kidney by approximately 40%, 24%, and 18%, respectively. The liver and kidney weights in Group VIII were comparable to those of the negative control Group I

**Table (5):** Effect of probiotics on liver, spleen, and kidney weight of the rats.

|  |  |  |  |
| --- | --- | --- | --- |
|  | **Liver (%)** | **Kidney (%)** | **Spleen (%)** |
| **Group I** | 4.50±0.35<sup>a</sup> | 0.65±0.21<sup>a</sup> | 0.53±0.09<sup>a</sup> |
| **Group II** | 8.30±0.89<sup>c</sup> | 0.85±0.38<sup>c</sup> | 1.03±0.12<sup>c</sup> |
| **Group III** | 6.40±0.44<sup>c</sup> | 0.79±0.13<sup>b</sup> | 0.84±0.06<sup>b</sup> |
| **Group IV** | 5.80±0.39<sup>b</sup> | 0.74±0.24<sup>b</sup> | 0.71±0.08<sup>b</sup> |
| **Group V** | 5.90±0.61<sup>b</sup> | 0.72±0.11<sup>b</sup> | 0.69±0.09<sup>b</sup> |
| **Group VI** | 5.8± 0.54<sup>b</sup> | 0.71± 0.22<sup>b</sup> | 0.69± 0.06<sup>b</sup> |
| **Group VII** | 5.9± 0.48<sup>b</sup> | 0.71± 0.31<sup>b</sup> | 0.67± 0.11<sup>b</sup> |
| **Group VIII** | 4.9± 0.58<sup>a</sup> | 0.70± 0.19<sup>a</sup> | 0.64± 0.09<sup>b</sup> |
| Data are presented as mean ± SD;<br>Means with different lowercase letters in the same column are significantly different (p<0.05). |  |  |  |

### 3.5. Effect of probiotics on the lipid profile of the rats

As expected, the positive control group (Group II) had a significantly worse lipid profile than the other groups (p<0.05) (Table 6). Total cholesterol (TC), triglycerides (TG), and low-density lipoprotein (LDL) levels were 2, 3, and 7 times higher, respectively, compared to Group I, while high-density lipoprotein (HDL) was nearly halved. Adding yogurt (T0) to the rats’ diet in Group III did not significantly affect their lipid profile compared to Group II (p > 0.05), as rats in Group III still exhibited high levels of TG, TC, and LDL, and low levels of HDL.

**Table (6):** Effect of probiotics on the lipid profile of the rats.

| **Table (6): Effect of probiotics on the lipid profile of the rats** |  |  |  |  |
| --- | --- | --- | --- | --- |
|  | **TC (mg/dl)** | **TG (mg/dl)** | **HDL (mg/dl)** | **LDL (mg/dl)** |
| **Group I** | 70.7±3.9<sup>a</sup> | 88.4±6.9<sup>a</sup> | 42.2±7.3<sup>a</sup> | 10.5±3.6<sup>a</sup> |
| **Group II** | 141.8±2.1<sup>c</sup> | 231.1±7.2<sup>c</sup> | 19.8±5.5<sup>c</sup> | 74.2±6.1<sup>c</sup> |
| **Group III** | 114.6±3.3<sup>c</sup> | 202.8±7.7<sup>c</sup> | 22.8±3.1<sup>c</sup> | 49.5±4.9<sup>c</sup> |
| **Group IV** | 92.4±2.9<sup>b</sup> | 170.7±7.1<sup>b</sup> | 31.5±4.4<sup>b</sup> | 27.9± 4.3<sup>b</sup> |
| **Group V** | 95.1±4.3<sup>b</sup> | 180.3±6.4<sup>b</sup> | 30.6±5.3<sup>b</sup> | 28.1±5.4<sup>b</sup> |
| **Group VI** | 94.8±3.5<sup>b</sup> | 171.8±7.5<sup>b</sup> | 32.9±7.9<sup>b</sup> | 29.3±2.1<sup>b</sup> |
| **Group VII** | 96.3±2.5<sup>b</sup> | 176.7±6.8<sup>b</sup> | 32.1±4.1<sup>b</sup> | 29.2±3.9<sup>b</sup> |
| **Group VIII** | 85.1±2.8<sup>b</sup> | 122.3±5.9<sup>b</sup> | 36.1±5.2<sup>a</sup> | 25.8±2.6<sup>b</sup> |
| Data are presented as mean ± SD;<br>Means with different lowercase letters in the same column are significantly different (p<0.05).<br>(TC) Cholesterol; (TG) Triglyceride; (HDL) High Density Lipoprotein; (LDL) Low Density Lipoprotein |  |  |  |  |

In contrast, the addition of probiotic-rich yogurt in Groups IV-VIII significantly improved the adverse effects of the HFD on the lipid profile of the rats (Table 6). Notably, yogurt with mixed probiotics (Group VIII) lowered TC, TG, and LDL by 40%, 47%, and 66%, respectively, while increasing HDL by 83% compared to Group II.

### 3.6. Effect of probiotics on the liver function of the rats

The HFD also significantly increased the ALT activity and reduced the albumin content in the positive control (Group II) by nearly 70% and 7%, respectively, compared to Group I (Table 7). Control yogurt (T0) in Group III ameliorated the adverse effect of the HFD on the hepatocytes; it significantly reduced the ALT activity to 44.60 U/L. Additionally, adding the probiotics to the yogurts attenuated the MASLD in the rats. Probiotic yogurt recovered the albumin in the plasma in groups IV-VIII by nearly 4-5%, with no significant differences between the probiotic strains. Besides, the probiotics exhibited a notable decrease in ALT activity compared to that in Group II, with ALT levels decreased by 13-28% in groups IV-VIII, and Group VIII showing the most pronounced effect.

**Table (7):** Effect of probiotics on the liver function of the rats.

| **Table (7): Effect of probiotics on the liver function of the rats** |  |  |
| --- | --- | --- |
|  | **ALT (U/L)** | **Albumin (mg/dl)** |
| **Group I** | 30.2± 2.1<sup>a</sup> | 3.37± 0.17<sup>a</sup> |
| **Group II** | 51.1±4.8<sup>c</sup> | 3±0.23<sup>b</sup> |
| **Group III** | 44.6±3.9<sup>b</sup> | 3.03±0.22<sup>b</sup> |
| **Group IV** | 41± 2.7<sup>b</sup> | 3.24±0.23<sup>a</sup> |
| **Group V** | 43.2± 3.3<sup>b</sup> | 3.22±0.22<sup>a</sup> |
| **Group VI** | 42.4± 4.2<sup>b</sup> | 3.23±0.19<sup>a</sup> |
| **Group VII** | 43.1±2.7<sup>b</sup> | 3.21±0.2<sup>a</sup> |
| **Group VIII** | 36.9±2.9<sup>b</sup> | 3.29±0.21<sup>a</sup> |
| Data are presented as mean ± SD;<br>Means with different lowercase letters in the same column are significantly different (p<0.05). |  |  |

### 3.7. Effect of probiotics on the glucose and insulin levels of the rats

The HFD significantly increased the glucose and the insulin in the plasma of the rats in Group II, reaching 52.10 mg/dl and 2.17 μIU/ml, respectively, compared to 19.60 mg/dl (glucose) and 0.69 μIU/ml (insulin) in Group I (Table 8). Including control yogurt in the rats’ diet did not significantly affect the glucose and insulin levels in the plasma compared to Group II (p>0.05). Interestingly, the probiotics significantly reduced glucose and insulin levels in the rats’ plasma. Notably, Group VIII, which received T5 probiotics yogurt, had the lowest glucose (28.60 mg/dl) and insulin (0.84 μIU/ml) levels.

**Table (8):** Effect of probiotics on the glucose and insulin levels of the rats.

|  |  |  |
| --- | --- | --- |
|  | **Glucose (mg/dl)** | **Insulin (μIU/ml)** |
| **Group I** | 19.6± 2.1<sup>a</sup> | 0.69± 0.11<sup>a</sup> |
| **Group II** | 52.10±3.00<sup>c</sup> | 2.17±0.19<sup>c</sup> |
| **Group III** | 41.80±3.40<sup>c</sup> | 1.45±0.08<sup>c</sup> |
| **Group IV** | 37.00±2.40<sup>b</sup> | 1.09±0.14<sup>b</sup> |
| **Group V** | 36.20±1.90<sup>b</sup> | 1.14±0.16<sup>b</sup> |
| **Group VI** | 31.10±2.80<sup>b</sup> | 1.04±0.10<sup>b</sup> |
| **Group VII** | 32.40±3.70<sup>b</sup> | 1.22±0.17<sup>b</sup> |
| **Group VIII** | 28.60±2.60<sup>b</sup> | 0.84±0.09<sup>a</sup> |
| Data are presented as mean ± SD;<br>Means with different lowercase letters in the same column are significantly different ($p<0.05$). | | |

### 3.8. Effect of probiotics on the Intestinal microbes

The microbial analysis of the rats’ intestines is shown in Table (9). Overall, the HFD increased the total coliform and staphylococci content while decreasing the total bacterial count in the intestine of the rats in Group II, compared to those on a normal chow diet (Group II). However, introducing T5 probiotic yogurt (containing a mix of probiotics) in Group VIII led to a significant decrease in coliform and staphylococci count, along with an increase in the total bacterial count, compared to Group II.

**Table (9):** Effect of probiotics on the microbial count of rat’s intestine.

| **Table (9): Effect of probiotics on the microbial count of rat's intestine** |  |  |  |
| --- | --- | --- | --- |
|  | **Total count (CFU/ml)** | **Coliform (CFU/ml)** | **Staphylococci (CFU/ml)** |
| **Group I** | 5.3×10<sup>7</sup> | 1.5×10<sup>6</sup> | 2.4×10<sup>8</sup> |
| **Group II** | 4.1×10<sup>7</sup> | 3.9×10<sup>6</sup> | 8.1×10<sup>8</sup> |
| **Group VIII** | 6.3×10<sup>7</sup> | 1.8×10<sup>6</sup> | 3.8×10<sup>8</sup> |

### 3.9. Effect of probiotics on the liver histology

In Group I, the liver section showed normal hepatocyte cords with healthy hepatic cells (black arrows) and a clear central vein (red arrows) **(Figure 2A; Table 10)**. In contrast, the HFD in Group II led to the loss of normal hepatocyte architecture. The liver cells exhibited rounded and pyknotic nuclei (black arrows), a focal area of hepatic necrosis with mononuclear cell infiltration (blue arrows), and large degenerated areas with centrilobular congestion, congested dilated portal tracts, and haemorrhage (red arrows) **(Figure 2B)**.

**Figure 2:**
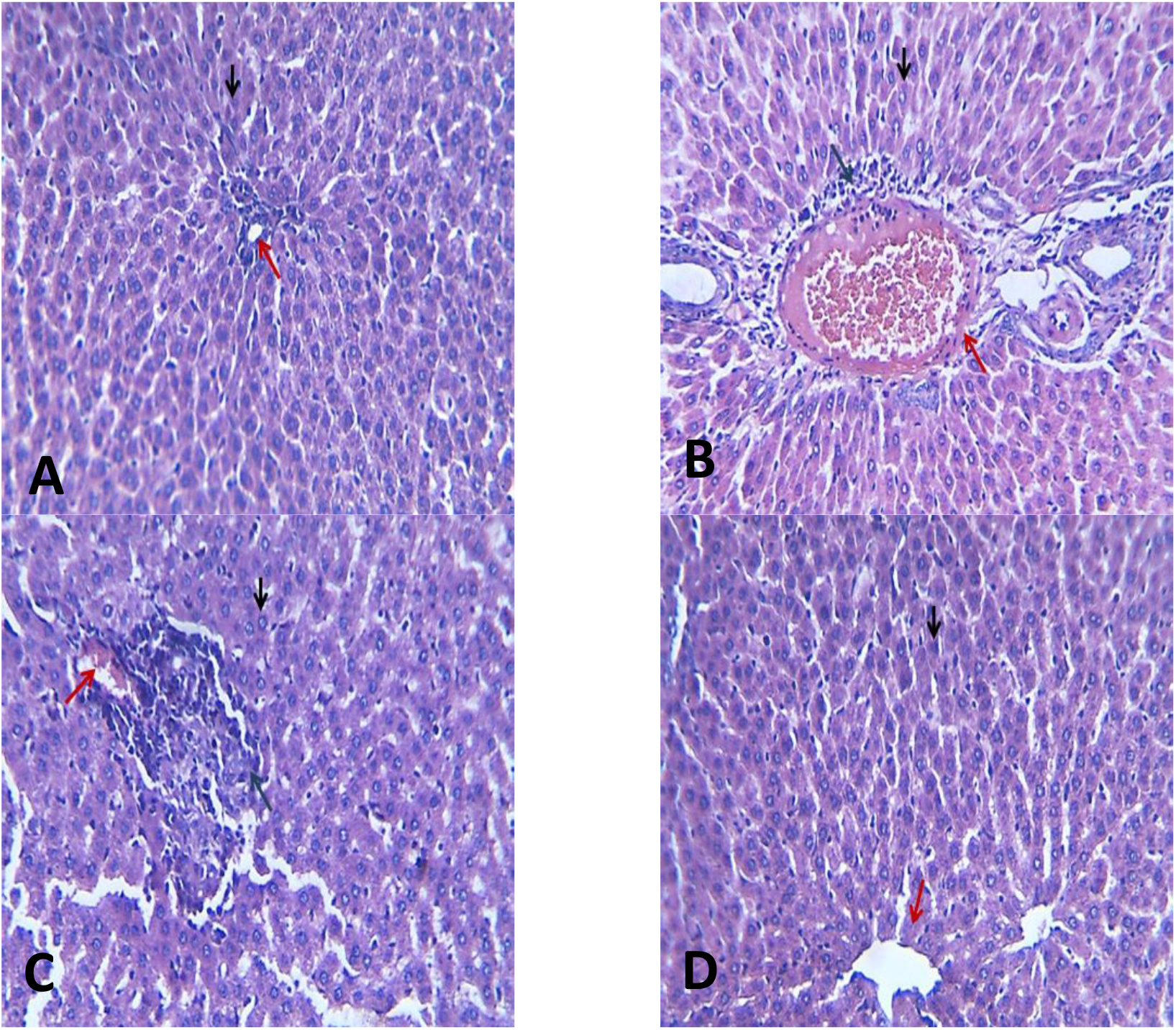
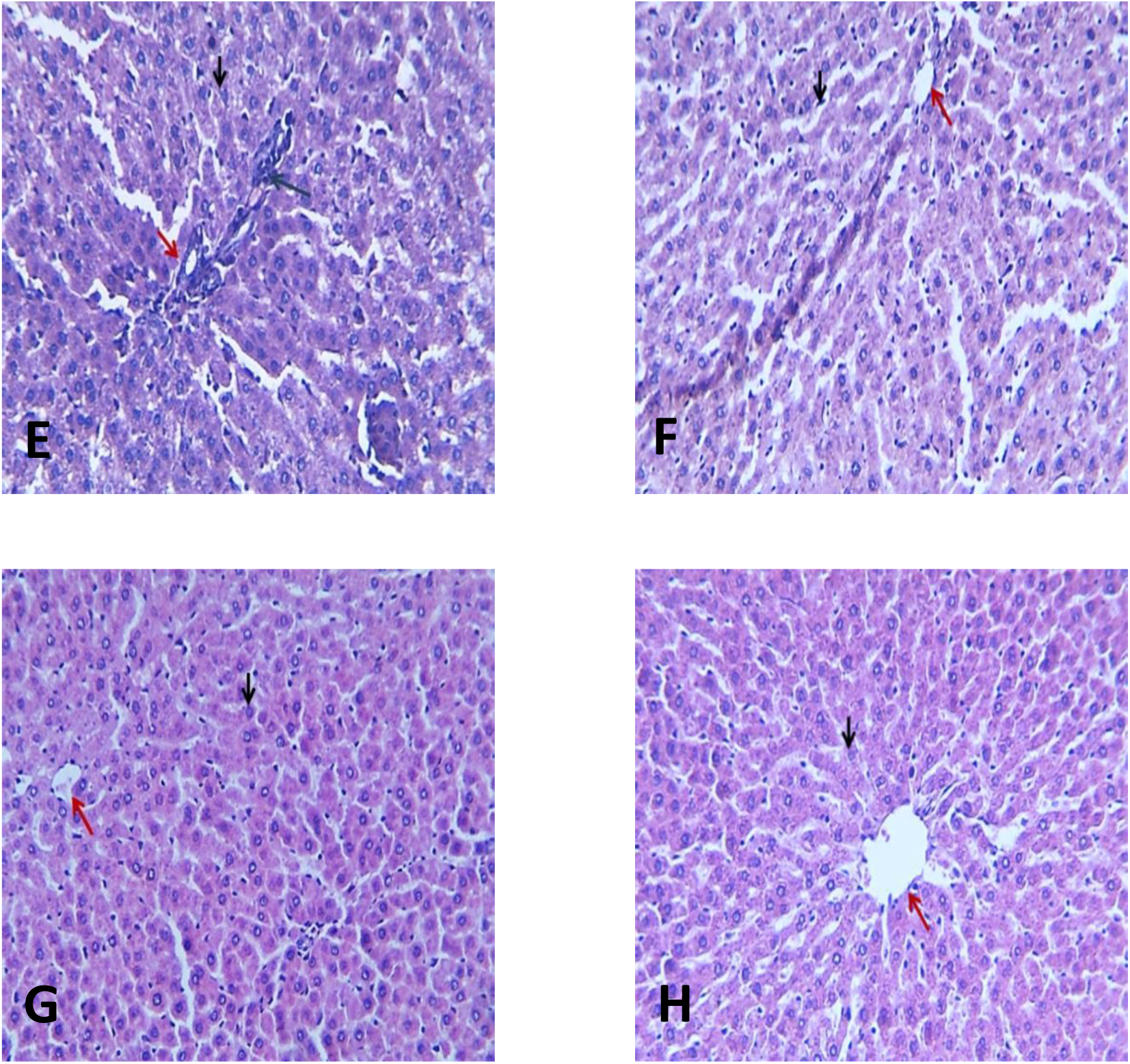
Effect of probiotic administration on liver histology.

**Table (10):**
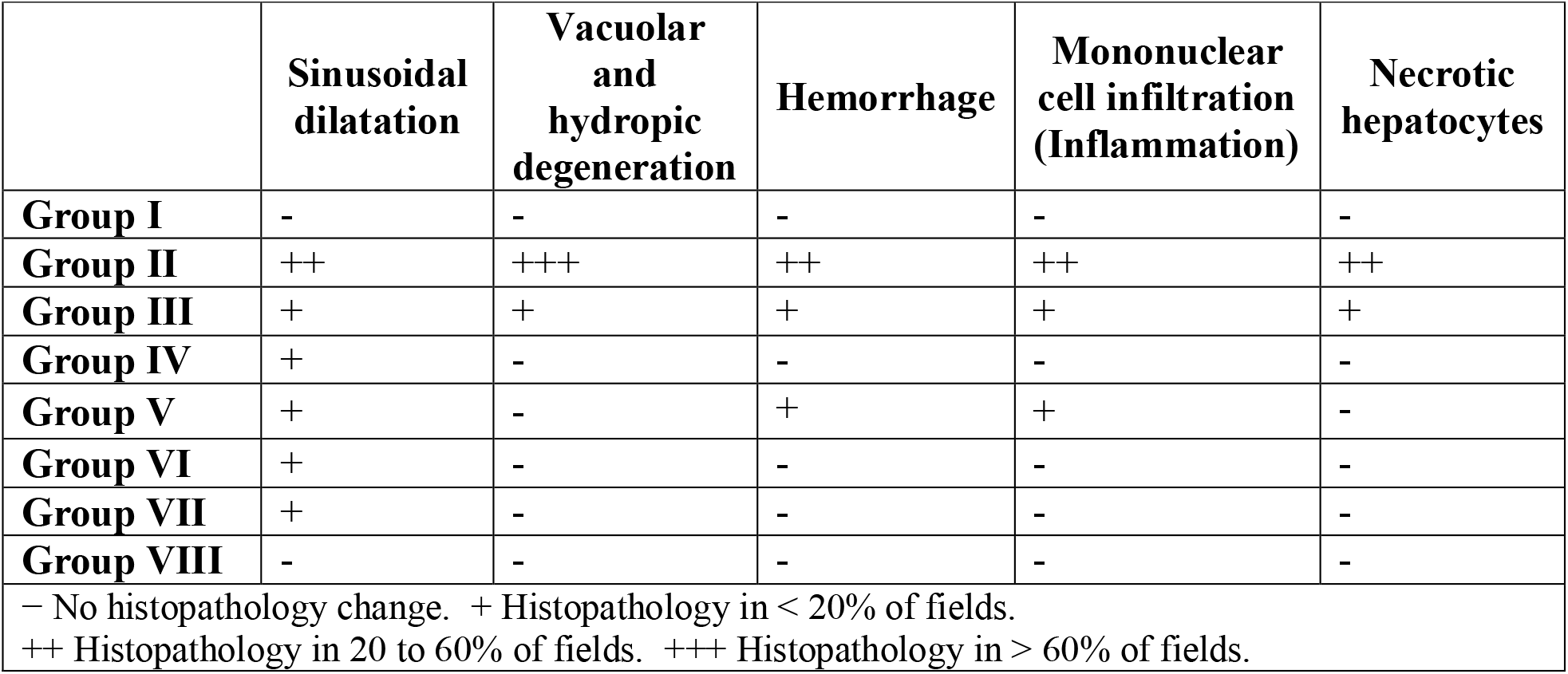
Effect of probiotic administration on liver histological scores.

Adding yogurt (T0) to the diet in Group III slightly mitigated the HFD’s adverse effects on the liver **(Figure 2C; Table 10)**. Although the liver section in Group III still showed a loss of normal hepatocyte architecture, there were fewer rounded and pyknotic nuclei (black arrows), smaller areas of focal hepatic necrosis (blue arrows), and reduced degeneration (red arrows) compared to Group II.

The effect of probiotics on the liver was strain-dependent. In Group V, *Lactobacillus plantarum* led to mostly normal hepatocytes (black arrows), with a small degenerated area (blue arrows) and a minor haemorrhagic region, along with a clear central vein (red arrows) **(Figure 2E)**. Meanwhile, *Lactobacillus rhamnosus* in Group IV **(Figure 2D)**, *Lactobacillus acidophilus* in Group VI **(Figure 2F)**, *Bifidobacterium lactis* in Group VII **(Figure 2G)**, and the mixed probiotic yogurt in Group VIII **(Figure 2H)** resulted in normal hepatocyte cords with healthy hepatic cells and active dilated sinusoids (black arrows) and clear central veins (red arrows).

## 4. Discussion

Probiotic-rich yogurt could mitigate the adverse effects of MASLD, particularly yogurt containing a mixture of probiotics. Yogurt enriched with *Lactococcus rhamnosus*, L*actobacillus plantarum, Lactobacillus acidophilus*, and *Bifidobacterium lactis* showed potential effects on HFD-induced MASLD. The probiotics efficiently reduced the weight of the organs, LDL, cholesterol, triglycerides, ALT activity, glucose, and insulin levels, while recovering the HDL and albumin levels. Moreover, the histological analysis revealed that probiotic-rich yogurt resulted in a normal hepatocyte structure. Compared to a single probiotic strain, the probiotic mixture had a more pronounced effect on the host’s health due to their synergistic and complementary actions within the gut. As a result, the host animals could show diverse signs of improved health and amelioration of metabolic syndromes (**Yoo, Kim et al. 2013)**.

In the present study, the high-fat diet (HFD) significantly increased the lipid profile (TG, TC, and LDL) of Group II compared to Group I, which was fed a normal chow diet. These findings align with the results of (Alnami, Bima et al.) and (Wang, Zhang et al.) The increase in lipid levels may be associated with the accumulation of fat droplets observed in the livers of the positive control rats (Group II), leading to the development of various degrees of MASLD, along with macro vascular and micro vascular steatosis. Diets such as HFD can change the conformation of gut microbiota by increasing gut permeability, reducing bacterial LPS removal, increasing levels of bacterial components, and inducing metabolic endotoxemia. Endotoxemia, in turn, may trigger subclinical inflammation, leading to numerous metabolic dysfunctions **(Kirpich, Marsano et al. 2015, Ojeda, Bobe et al. 2016, Chakaroun, Massier et al. 2020)** Consequently, fat metabolism may be disrupted, increasing free fatty acid uptake, triglyceride lipogenesis, and fat deposition in the hepatic tissues. Besides, a bacterial imbalance can take place, which can contribute to the loss of gut barrier function, leading to bacterial translocation and induction of inflammatory responses in the liver **(Sabate, Jouet et al. 2008, Jiang, Wu et al. 2015, Briskey, Heritage et al. 2016)**. Ingestion of a high-fat diet alters the intestinal microbiota and increases gut-derived inflammatory agents by renewing the bowel flora in conditions of high-fat diet-induced steatosis. The progression of inflammation is also an important factor, as it is associated with the development of obesity and MASLD, resulting from changes in diet and microbiota **(de La Serre, Ellis et al. 2010)**. High-fat diets, in particular, can lead to endotoxemia due to elevated levels of bacterial LPS in the blood **(Laugerette, Vors et al. 2020)**.

Herein, the HFD significantly increased liver weight in all HFD-fed groups (Groups II-VIII) compared to Group I, which was fed a chow group. The obtained findings are consistent with those of (Helal, El-Wahab et al.), who observed enlarged and ballooned hepatocytes in fatty liver-induced rats, with macro-vacuoles scattered throughout the cytoplasm of cells across the hepatic lobule. Groups II-VIII showed significant fat deposition within hepatocytes compared to Group I. However, the group receiving the probiotic mixture (Group VIII) showed a significant reduction in liver weight compared to the positive control (Group II). These results align with the findings of (Rishi, Arora et al.) who reported that probiotic administration improved the liver morphology.

The TG and TC levels were elevated in all HFD-fed groups, with Group II showing the highest levels. These findings agree with previous literature (Liu, Han et al.), (Lee, Chiang et al.) and (Hussain, Cho et al.). The results also showed a decrease in high-density lipoprotein (HDL) and an increase in low-density lipoprotein (LDL) in the positive control group. Chronic intake of the fat-enriched diet could induce fatty liver, resulting in elevated LDL and reduced HDL levels **(Sigrist-Flores, Ponciano-Gomez et al. 2019)**, which are linked to hepatic steatosis **(Briseño-Bass, Chávez-Pérez et al. 2019)** and MASLD **(DeFilippis, Blaha et al. 2013)**. On the other hand, the probiotic mixture in Group VIII significantly lowered the TG and TC levels compared to the positive control. Similarly, probiotic supplementation dramatically decreased total cholesterol and triglycerides **(Kullisaar, Zilmer et al. 2016)**. Probiotics are thought to decrease cholesterol by de-conjugating bile salt, blocking the reabsorption and promoting their subsequent excretion, thus reducing cholesterol recycling (**Begley, Hill et al. 2006)**. The probiotic mixture also showed a pronounced effect in mitigating the adverse effects of the HFD in Group VIII. It significantly increased the HDL levels and decreased the LDL levels. The reduction of the LDL could be attributed to the probiotic-induced recovery of liver low-density lipoprotein receptor (LDLr), facilitating the liver to absorb plasma LDL **(Song, Park et al. 2015**).

ALT is considered a crucial non-invasive marker of inflammation **(Liu, Que et al. 2014**). Rats fed on HFD showed elevated ALT activity levels compared to those on a chow diet. This could be a consequence of increased lipogenesis, lipids accumulation in the hepatic cells, and focal periportal inflammation. In contrast, the ALT level decreased in Group VIII, which received a probiotic mixture. This result aligns with previous studies of (ADESIJI, OWOLABI et al.) and (Li, Shi et al.) who reported that probiotics can decrease liver enzyme levels. Likewise (Manzhalii, Virchenko et al.) reported that patients who was receiving probiotics have shown a notable reduction in ALT levels, hepatic inflammation, and liver stiffness.

The results also revealed a significant decrease in albumin levels in the positive control group compared to the other groups. This finding agrees with previous studies of (Grgurevic, Podrug et al.) and (Kawaguchi, Sakai et al.) who found that the patients who had MALSD showed a low level of serum albumin. The probiotic mixture in Group VIII significantly recovered the albumin levels in the rats’ serum. These obtained results are consistent with previous literature (Ayyat, Al-Sagheer et al.) who reported that probiotics can significantly increase serum total protein and albumin levels. However, (Alkhalf, Alhaj et al.) stated that probiotic supplementation had no effect on the serum total protein or albumin. The discrepancies between the results could be attributed to various factors, such as the probiotic strain and dose; experimental duration and conditions; host’s type and age.

Furthermore, serum glucose and insulin are common markers of MASLD. The experiment showed that the HFD raised the glucose levels in Group II compared to the other groups these results aligns with (Helal, El-Wahab et al.) and (Cho, Namgung et al.) who found that high glucose levels have been reported as a sign of fatty liver Interestingly, Group VIII, which received the probiotic mixture, showed a significant lower glucose level. This finding aligns with previous research of (ADESIJI, OWOLABI et al.) which shows a reduced glucose levels in probiotic-treated rats. This observation may be attributed to the appropriate insulin release, which helps to control blood glucose levels (**Andreasen, Larsen et al. 2010)**. Probiotics may enhance endogenous insulin production by promoting glucose storage in the liver, increasing the body’s glucose utilization, or improving declining beta-cell activity (**Duan, Liu et al. 2015)**.

The experiment showed that the HFD induced an increase in MASLD markers, such as liver function markers, lipid profile, glucose, insulin resistance, final body weights, and body weight gains. In contrast, the administration of a probiotic mixture could counter these changes, promoting overall health. Previous studies of (Bloemendaal, Szopinska-Tokov et al.) and (Liu, Que et al.) have shown that various can modulate gut bacterial structure and composition, especially when disturbed by a high-fat diet. Finally, MASLD can be effectively treated using probiotics to regulate the gut-liver axis. In this study, a mixture of *Bifidobacterium* and *Lactobacillus* strains affected host inflammation and intestinal microbial composition, consistent with the findings of previous research of (Hong, Kim et al.), (Mazloom, Siddiqi et al.), (da Silva, Casarotti et al.) and (Ashaolu and Fernandez-Tome).Treatment of MASLD with different probiotic mixtures induced modifications within the intestinal microbiota, which help attenuate metabolic disruptions by reducing serum lipid profiles and inflammatory biomarkers.

## 5. Conclusion

In conclusion, the study highlights the pronounced effect of the probiotic mixture in attenuating liver steatosis and hepatic inflammation by regulating the host metabolism via modulation of the gut micro biome, specifically in terms of gut microbial composition. Altogether, these findings would be of great value for future application of the probiotic mixture as preventive or early-stage therapeutics in MASLD treatment. The probiotic mixture showed a clear ability to improve the blood biochemical markets of rats fed with HFD, recovering the cholesterol, triglycerides, HDL, LDL, glucose, insulin, liver weight, and body weight closer to the levels observed in the control group (fed a normal diet). Besides, the mixture significantly modulated the intestinal microbiota by reducing the coliform and *staphylococcus aurues* counts

The current data suggest that by retarding and reversing dysbiosis, probiotics can improve hepatic inflammation, histology, and function as reflected in biochemical markers and liver biopsy samples. Theoretically, probiotics can be used in combinations as MASLD-targeted therapies, given that probiotics are safe, inexpensive, and have no known adverse effects with long-term use. Therefore, probiotics supplementation seems to be a practical therapeutic strategy in MASLD/MASH management.

## Declarations

### Ethics approval

All experiments were performed according to relevant guidelines and regulations. The study has been approved by the Ethics Committee of Alexandria University, Egypt **(ALEXU-IACUC) (AU: 19/22/10/20/3/24)**

### Consent to participate

Not applicable.

### Data Availability Statement

All data generated and analyzed in this study are included in this article.

### Competing interests

The authors have declared no conflict of interest.

### Funding

This research was not supported by any fund.

## Notes

### Competing Interest Statement

The authors have declared no competing interest.

